# Rewiring c-Myc Transcriptional Activity with an O-GlcNAcylation Targeting Chimera (OGTAC)

**DOI:** 10.64898/2026.05.04.722559

**Authors:** Tongyang Xu, Zhihao Guo, Khadija S. Khan, Yunpeng Huang, Bowen Ma, Jialin Liu, Dean W. Felsher, Billy Wai-Lung Ng

## Abstract

c-Myc is a transcription factor that drives tumorigenesis in many cancers. It is notoriously difficult to directly target c-Myc, mainly due to its lack of well-defined druggable pockets. O-linked β-N-acetylglucosamine modification (O-GlcNAcylation) is a post-translational modification (PTM) playing an important role in regulating c-Myc functions in cancer. However, previous studies have primarily relied on global perturbations to investigate c-Myc O-GlcNAcylation, making it difficult to determine its direct functional consequences due to concurrent cellular effects. Here, we report a bifunctional O-GlcNAcylation TArgeting Chimera (OGTAC) molecule, which can induce the proximity of c-Myc and O-GlcNAc transferase (OGT) in living cells, thereby enhancing the O-GlcNAcylation of c-Myc. The c-Myc-targeting OGTAC exhibits anti-proliferation effect against cancer cells. Mapping of c-Myc occupancy on genome indicates that OGTAC rewires c-Myc transcriptional activity and reprograms expression of the downstream oncogene *MALAT1*, in an O-GlcNAcylation-dependent manner. Overall, OGTAC presents a novel chemically induced proximity (CIP)-based tool to target and rewire c-Myc activity in cancer.

**Graphic abstract:** 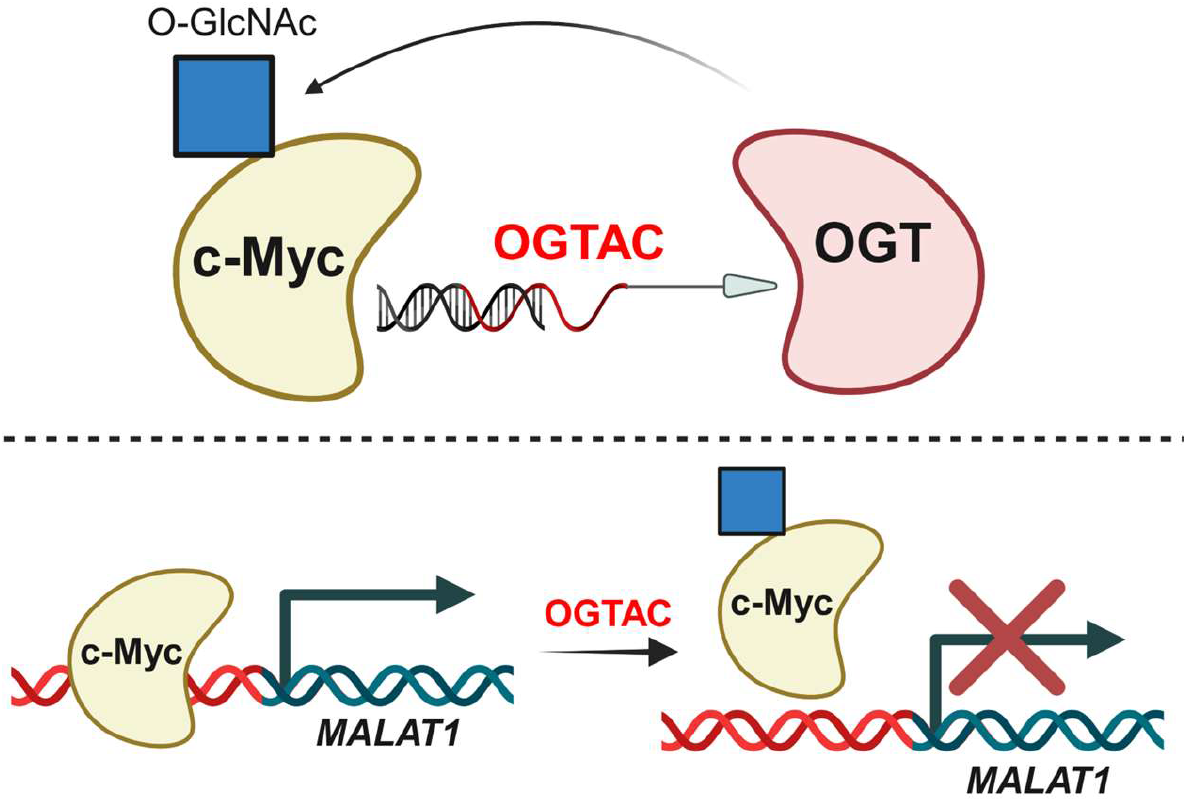

## Introduction

The oncoprotein c-Myc is an essential transcription factor, encoded by the oncogene *MYC*. Its dysregulation is a hallmark of tumor progression across diverse cancer types.^1-4^ This central role renders c-Myc a compelling target in cancer. However, its lack of well-defined binding pockets hinders drug development. Whereas previous studies primarily focused on developing c-Myc inhibitors and degraders,^5, 6^ the present study proposes using an O-GlcNAcylation Targeting Chimera (OGTAC) to target c-Myc O-GlcNAcylation and modulate c-Myc transcriptional activity. OGTAC represents a heterobifunctional small molecule that enables protein-specific O-GlcNAcylation in living cells through induced proximity to O-GlcNAc transferase (OGT).^7-9^

O-GlcNAcylation is important for functions of transcription factors.^10^ Regarding c-Myc, previous studies have primarily regulated its O-GlcNAcylation through inhibition or knockdown of OGT or O-GlcNAcase (OGA).^11-14^ O-GlcNAcylation of c-Myc at Thr58 and Ser415 has been reported to affect its phosphorylation, stability, subcellular localization, and apoptosis and other downstream processes across multiple cancer types.^11-17^ The present study aims to specifically target c-Myc using OGTAC to elicit direct functional effects, thereby avoiding perturbation of cellular O-GlcNAcylation patterns induced by prior methods (Figure 1). Overall, this proof-of-concept study presents OGTAC as an alternative means modulating c-Myc transcriptional activities.

**Figure 1.**
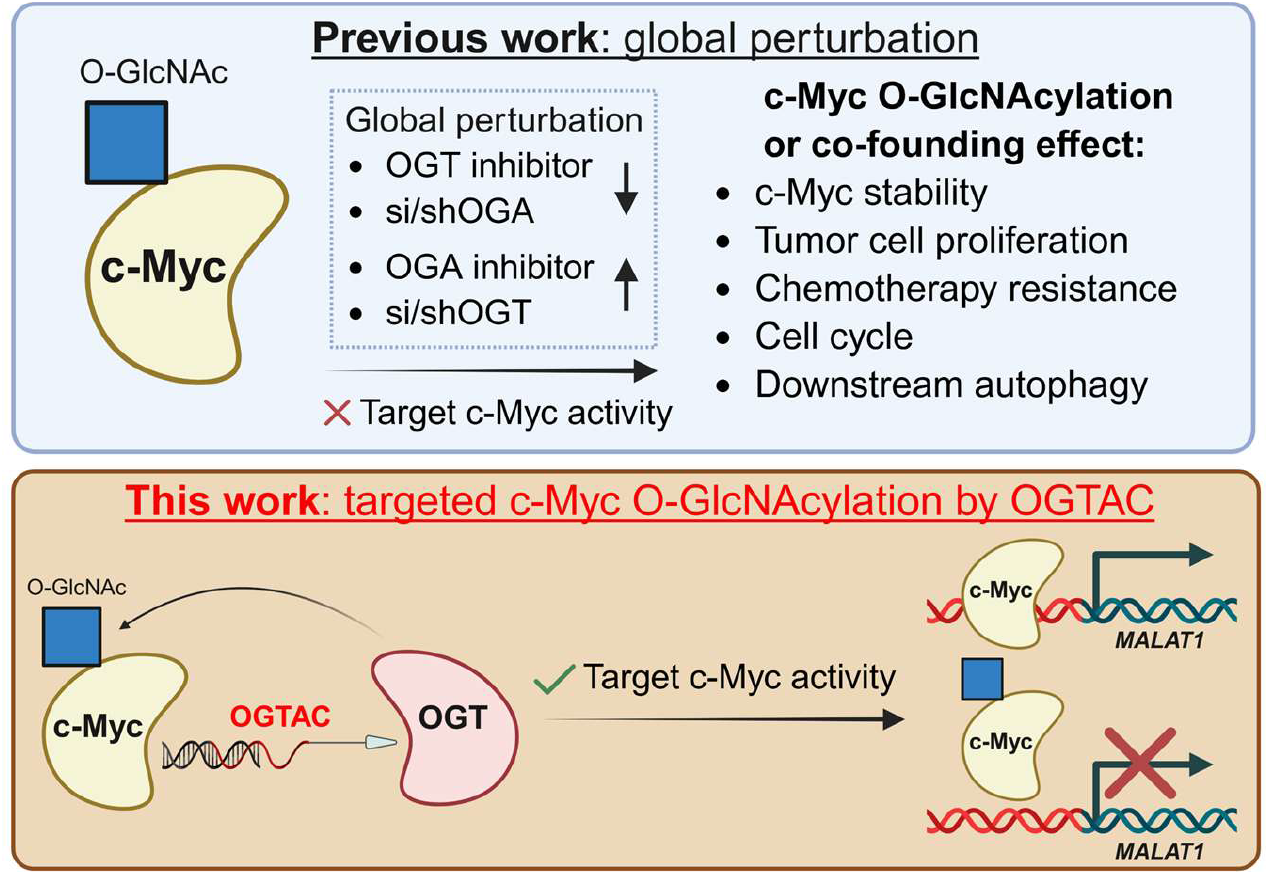
Schematic presentation of previous approaches that globally regulate O-GlcNAcylation, and the use of OGTAC to specifically target c-Myc O-GlcNAcylation.

## Results and Discussions

### Design, Synthesis, and Evaluation of the c-Myc-Targeting Bifunctional Molecules

To target c-Myc, we designed an OGTAC that combines a reported extended E-box oligonucleotide (oligo) for c-Myc binding,^18^ with AP1867 to recruit FKBP12^F36V^-fused OGT (fOGT) (Figure 1, 2A; Supplementary Table 1).^7^ Previous studies have successfully regulated the stability of c-Myc and other transcription factors using this class of oligonucleotide warheads; importantly, these warheads are compatible with *in vivo* studies.^18-22^ The recruitment of c-Myc by a biotin-conjugated oligo was first evaluated (Supplementary Table 1; Supplementary Figure 1A). Biotin pull-down assay demonstrated dose-dependent enrichment of c-Myc from cell lysates by the biotin-conjugated oligo, whereas the biotin-conjugated scrambled DNA and the unbiotinylated oligo did not enrich c-Myc. Subsequently, the bifunctional oligo(5′ or 3′)-(PEG)_3_-AP1867 were synthesized via strain-promoted azide–alkyne cycloaddition (SPAAC)^23^ between oligo(5′ or 3′)-N_3_ and dibenzocyclooctyne-(PEG)_3_-AP1867 (DBCO-(PEG)_3_-AP1867), and were characterized by native PAGE, which revealed a 1,155 Da increase in apparent molecular weight after the click reaction (Figure 2A; Supplementary Figure 1B). Oligo lacking N_3_ modification, and the scrambled DNA, were subjected to the same reaction conditions with DBCO-(PEG)_3_-AP1867 and were used as negative controls, namely c-Myc-binding DNA complex (CBD, binding c-Myc but not recruiting OGT) and scrambled DNA complex (SD, neither binding c-Myc nor recruiting OGT) (Supplementary Table 1).

**Figure 2.**
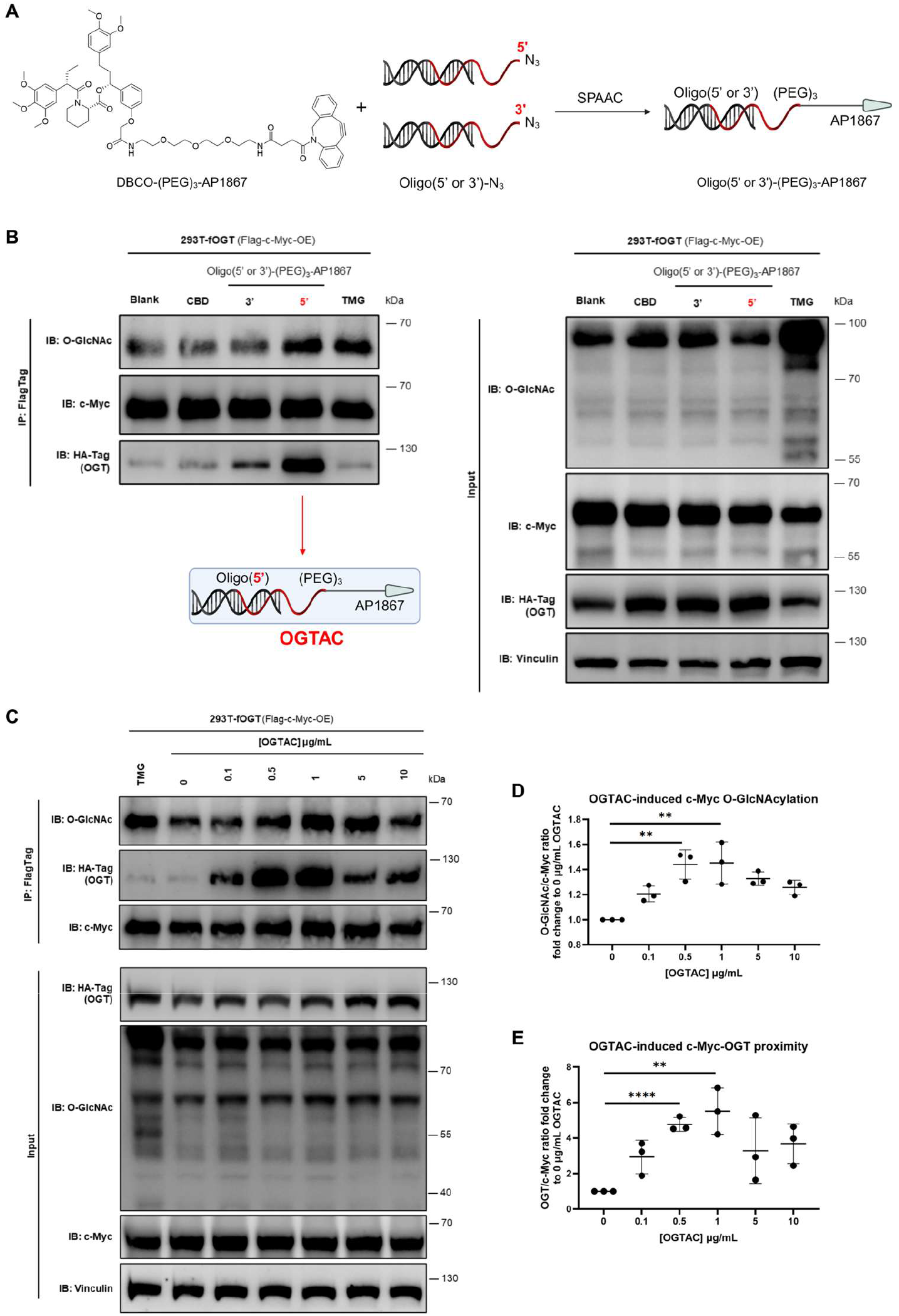
Design and evaluation of the c-Myc-targeting OGTAC. (A) Design of the c-Myc-targeting OGTAC, composed of the c-Myc-binding extended E-Box oligonucleotide (oligo) and the FKBP12^F36V^-binding AP1867. We synthesized the bifunctional molecules via a strain-promoted azide–alkyne cycloaddition (SPAAC) reaction between oligonucleotide (5′ or 3′)-N_3_ and DBCO-(PEG)_3_-AP1867. (B) Oligo(5’)-(PEG)_3_-AP1867 produced a stronger induction than the 3′-linked version and was therefore designated OGTAC. (C) Monitoring of c-Myc O-GlcNAcylation and c-Myc-OGT co-IP following dose-dependent OGTAC treatment. OGTAC at 1 μg/mL produced the maximal induction, and a hook effect was observed at higher concentrations. Thiamet-G (TMG) was used as the positive control. (D)Quantification of relative c-Myc O-GlcNAcylation (O-GlcNAc signal in IP normalized to c-Myc in IP), expressed as fold change relative to 0 μg/mL OGTAC. (E) Quantification of relative c-Myc-OGT co-IP (OGT signal in IP normalized to c-Myc in IP), expressed as fold change relative to 0 μg/mL OGTAC. Error bars represent the mean (SD) from *n* = 3 biologically independent experiments. Statistical significance for the quantifications was determined using an unpaired Student’s *t*-test. \*\**p* < 0.01; \*\*\*\**p* < 0.0001.

### OGTAC Induces c-Myc O-GlcNAcylation and c-Myc-OGT Proximity in Living Cells

O-GlcNAcylation of c-Myc was detected by immunoprecipitation (IP) and western blot (WB) (Supplementary Figure 2). To assess whether oligo-(PEG)_3_-AP1867 constructs can induce c-Myc O-GlcNAcylation and co-IP with OGT, 1 μg/mL of the oligo(5′ or 3′)-(PEG)_3_-AP1867 constructs or controls were applied to lysates of HEK293T cells co-overexpressing fOGT and Flag-c-Myc. Both oligo(5′)- and oligo(3′)-linked (PEG)_3_-AP1867 constructs induced O-GlcNAcylation, with the double-stranded duplex outperforming the unannealed single-stranded constructs (Supplementary Figure 3). To apply in living cells, both HEK293T and HeLa cell lines stably overexpressing fOGT (HEK293T-fOGT and HeLa-fOGT) were generated. HEK293T-fOGT cells overexpressing Flag-c-Myc were transfected with 1 μg/mL of the oligo(5′ or 3′)-(PEG)_3_-AP1867 or with CBD for 24 h. Only the oligo(5’)-(PEG)_3_-AP1867 produced the strongest induction (Figure 2B) and was designated OGTAC for subsequent experiments.

Next, a dose-dependent IP-WB analysis was performed to determine the optimal working concentration of OGTAC (Figure 2C-E). After 24 h treatment, 1 μg/mL OGTAC produced the greatest OGT engagement (∼6-fold) and specifically increased c-Myc O-GlcNAcylation by ∼1.5-fold (Figure 2C-E). The magnitude of c-Myc O-GlcNAcylation induced by OGTAC was comparable to that induced by the global O-GlcNAcylation inducer thiamet-G (TMG),^24^ whereas global O-GlcNAcylation levels were not altered by OGTAC (Figure 2B, C). Both co-IP and O-GlcNAcylation induction declined at 5 μg/mL, indicating a hook effect at this concentration. Furthermore, application of 1 μg/mL OGTAC to HeLa-fOGT cells for 24 h resulted in an over 2-fold increase in co-IP between endogenous c-Myc and OGT (Figure 3A, B). Immunofluorescence (IF) imaging of Cy3-CBD or Cy3-OGTAC and c-Myc indicated stronger colocalization in nucleus than that observed with Cy3-SD and c-Myc (Figure 3C, D; Supplementary Table 1). These results further support target engagement by OGTAC. While previous studies showed that O-GlcNAcylation of c-Myc mainly led to its stabilization,^11-14^ c-Myc under OGTAC treatment did not present significantly different stability compared to CBD, yet its destabilization was observed on c-Myc-oligo binding over the SD control (Supplementary Figure 4).

**Figure 3.**
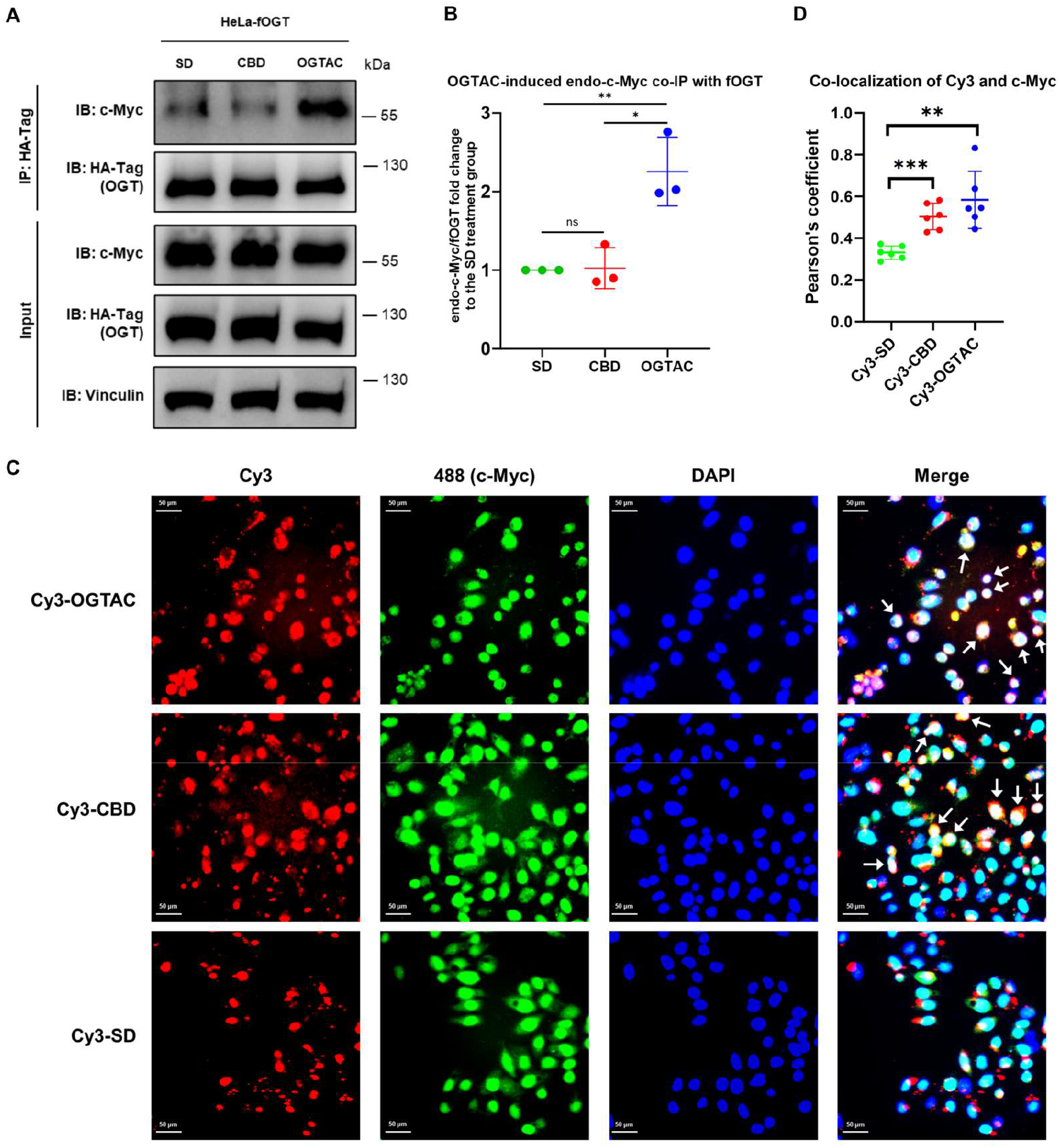
OGTAC engages and colocalizes with endogenous c-Myc in the nucleus. (A) 1 μg/mL OGTAC treatment for 24 h induced co-IP of endogenous c-Myc with OGT. (B)Quantification of relative endogenous c-Myc-OGT co-IP (c-Myc signal in IP normalized to OGT signal in IP), expressed as fold change relative to the SD control. (C)IF imaging of Cy3-conjugated constructs (red: Cy3-SD, Cy3-CBD, Cy3-OGTAC), c-Myc (green: 488), and DAPI (blue: nucleus), and merged channels. Cy3-OGTAC and Cy3-CBD, but not Cy3-SD, exhibited strong colocalization (white regions in the merged image, arrows) with endogenous c-Myc and the nucleus. Scale bar = 50 μm. (D)Quantification of Pearson’s correlation coefficient for colocalization between Cy3-conjugated constructs (Cy3-SD, Cy3-CBD, Cy3-OGTAC) and endogenous c-Myc (488), expressed as fold change relative to the Cy3-SD control. Error bars in (B) represent the mean (SD) from *n* = 3 biologically independent experiments. Error bars in (D) represent the mean (SD) from *n* = 6 biologically independent experiments. Statistical significance for the quantifications was determined using an unpaired Student’s *t*-test. \**p* < 0.1; \*\**p* < 0.01; \*\*\**p* < 0.001; ns, not significant.

Given OGTAC’s potency in inducing proximity and targeted c-Myc O-GlcNAcylation, the effect on cancer cell viability was subsequently investigated. HeLa-fOGT cells were treated with OGTAC at 0.1 - 5 μg/mL for 24 h, and a significant decrease in cell viability was observed relative to CBD and SD controls (Figure 4A). Notably, the concentration at which OGTAC suppressed viability corresponded to that producing maximal induction in the above assay (Figure 2C-E).

**Figure 4.**
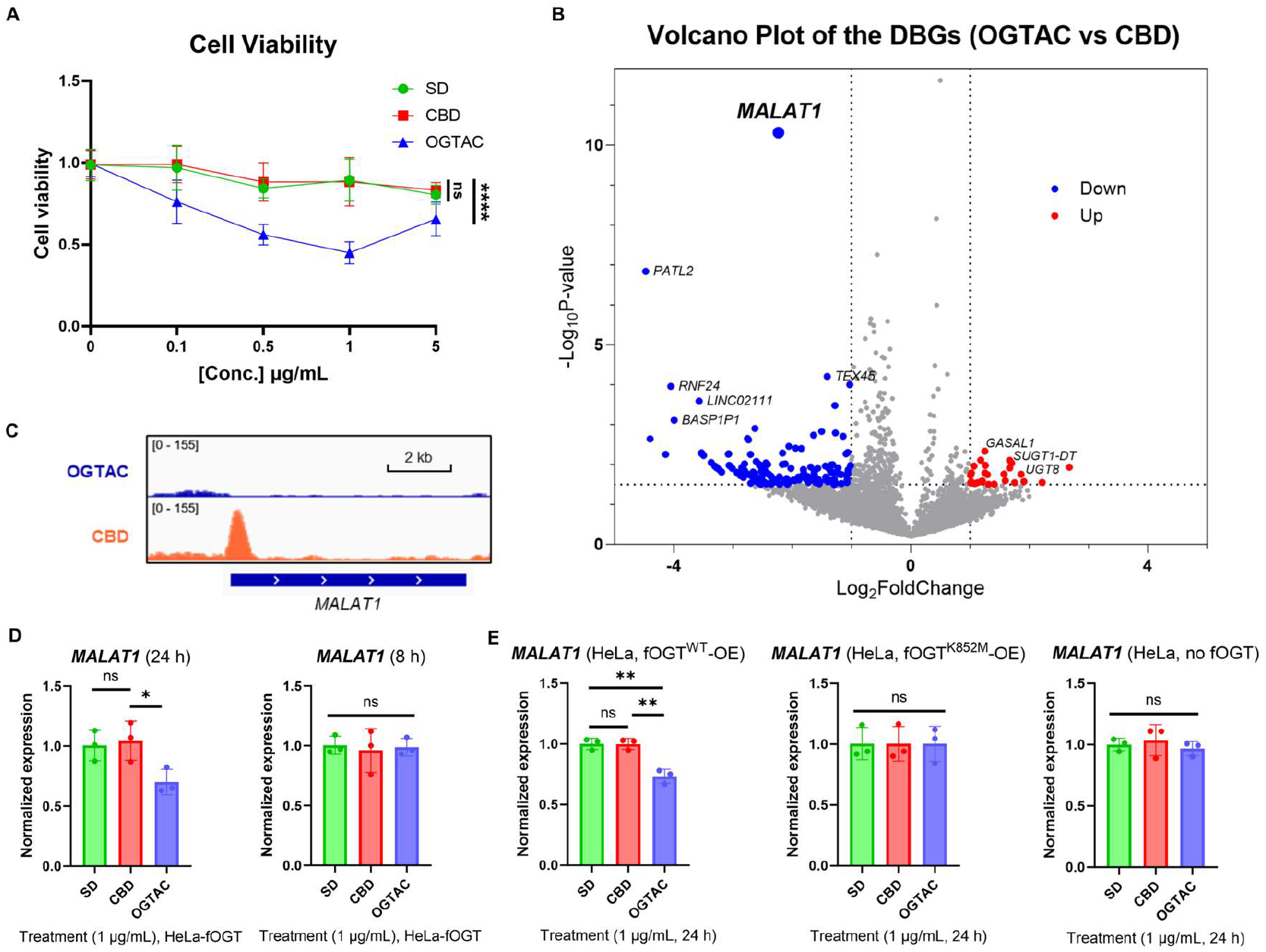
OGTAC inhibited HeLa viability and rewired c-Myc transcriptional activity. (A) CCK-8 assay showing reduced viability of HeLa-fOGT cells following OGTAC treatment relative to SD and CBD controls. Error bars represent the mean (SD) from *n* = 3 biologically independent experiments, each containing *n* = 3 technical replicates. Statistical significance was assessed using a two-way ANOVA. \*\*\*\**p* < 0.0001; ns, not significant. (B)Volcano plot of differentially bound genes (DBGs) of c-Myc identified by Cut&Tag-seq (OGTAC vs CBD). Results are from biological duplicates. A read-count threshold of 20 was applied to filter low-quality peaks, yielding 10,628 DBGs. (C)IGV genome-browser view of c-Myc binding at the *MALAT1* locus following OGTAC (blue) and CBD (orange) treatment. (D)Normalized expression of *MALAT1* and *MYC* in HeLa-fOGT cells after 8 h and 24 h treatment with 1 μg/mL OGTAC, expressed relative to SD and CBD controls. (E)Normalized expression of *MALAT1* and *MYC* in HeLa cells overexpressing fOGT^WT^, fOGT^K852M^, or vector control after 24 h treatment with 1 μg/mL OGTAC, expressed relative to SD and CBD controls. In panel (D) and (E), error bars represent the mean (SD) from *n* = 3 biologically independent experiments, where each point represents the mean of *n* = 3 technical replicates. Relative expression was calculated as target gene/HPRT1 and is presented as fold change relative to the SD control. Statistical significance for the quantifications was determined using an unpaired Student’s *t*-test. \**p* < 0.1; \*\**p* < 0.01; ns, not significant.

### OGTAC Rewires c-Myc Transcriptional Activity

To further delineate the mechanism by which the c-Myc-targeting OGTAC affects cell viability, Cleavage Under Targets and Tagmentation followed by sequencing (Cut&Tag-seq) of c-Myc was performed to monitor its genomic occupancy after 24 h treatment with 1 μg/mL OGTAC.^25^ Compared with the CBD control, OGTAC reconfigured the global genomic occupancy of c-Myc (Supplementary Figure 5A-C). A volcano plot of differentially bound genes (DBGs) indicated that OGTAC altered the c-Myc occupancy profile, with the majority of DBGs exhibiting decreased c-Myc binding (Figure 4B). Among DBGs, OGTAC decreased c-Myc occupancy at the *MALAT1* locus by over 4-fold relative to CBD treatment. Genome-browser views of Cut&Tag-seq data confirmed that OGTAC treatment resulted in near-complete loss of c-Myc occupancy at the *MALAT1* promoter (Figure 4C). *MALAT1* is a highly conserved long noncoding RNA that is frequently dysregulated and overexpressed in cancer.^26, 27^ c-Myc ChIP-seq data from the Encyclopedia of DNA Elements (ENCODE)^28^ suggests a strong c-Myc occupancy at the *MALAT1* promoter in cancer cell lines (Supplementary Figure 5D). We also found that c-Myc overexpression activated *MALAT1* transcription (Supplementary Figure 6A), which is consistent with previous studies.^19^ This relationship has been reported to facilitate c-Myc oncogenic activity, thereby forming a reciprocal regulatory loop.^29, 30^ RT-qPCR validation revealed that 24 h OGTAC treatment at 1 μg/mL significantly decreased *MALAT1* expression in both HeLa-fOGT and c-Myc-overexpressing HEK293T-fOGT cells relative to SD and CBD controls, whereas no such effect was observed at 8 h treatment (Figure 4D; Supplementary Figure 6B). Notably, *MYC* expression was also reduced by OGTAC in HeLa-fOGT cells, which may reflect feedback resulted from the downregulation of *MALAT1* (Supplementary Figure 6C).

### OGTAC Suppressed *MALAT1* Transcription in a c-Myc O-GlcNAcylation-Dependent Manner

Given that OGTAC treatment downregulated *MALAT1* expression, we subsequently investigated whether it was mediated by OGTAC-induced c-Myc O-GlcNAcylation or by induced c-Myc-OGT proximity. A catalytically inactive fOGT mutant (K852M)^31^ was generated, which lacked O-GlcNAc transferase activity both globally and toward c-Myc, but retained normal interaction with c-Myc (Supplementary Figure 7). RT-qPCR demonstrated that OGTAC decreased *MALAT1* expression only in the presence of wild-type fOGT (fOGT^WT^), but not with fOGT^K852M^ or vector control (Figure 4E). These results indicated that the effect of OGTAC on *MALAT1* is dependent on the targeted O-GlcNAcylation of c-Myc. It is noteworthy that O-GlcNAcylation of c-Myc with T58A and S415A (the previously reported c-Myc O-GlcNAcylation sites)^11, 16^ mutations was still induced by fOGT^WT^, suggesting additional c-Myc O-GlcNAcylation sites (Supplementary Figure 7).

## Conclusions

In summary, we present the c-Myc-targeting OGTAC by conjugating c-Myc-binding oligo with fOGT small molecule binder. This OGTAC constitutes the first reported tool to induce targeted c-Myc O-GlcNAcylation and thereby rewire c-Myc transcriptional regulation of downstream genes. Specifically, targeted O-GlcNAcylation of c-Myc by OGTAC downregulated *MALAT1* transcription and expression and may consequently impair cancer cell viability. OGTAC enables specific O-GlcNAcylation of c-Myc, avoiding broad proteome perturbation. Its modular, oligo-guided design permits adaptation to other transcription factors. Overall, as a novel approach that modulates c-Myc activity via targeted PTM rather than protein degradation, OGTAC has potential to be further developed as an anti-tumor tool in c-Myc-directed cancers.

## Acknowledgments

B.W.-L.N. acknowledges funding support from CUHK (CU Medicine Faculty Innovation Award; FIA2020/A/03), Guangdong-Hong Kong-Macao Joint Laboratory for New Drug Screening, Hong Kong RGC (General Research Fund; 14308724), the Peter Hung Pain Research Institute (PHPRI) Research Fund, and the Gerald Choa Neuroscience Institute (GCNI) Research Fund. The authors sincerely acknowledge Dr. Anja Deutzmann (Stanford University), Prof. Linda S. Gu (CUHK), Prof. Yvonne N.-W. Kam (CUHK), and Dr. Yalun Xie (CUHK) for scientific discussions.

