## Supporting Information for "Rewiring c-Myc Transcriptional Activity with an O-GlcNAcylation Targeting Chimera (OGTAC)"

### *Methods and Materials.*

#### Cell culture, DNA constructs and reagents

Experiments were performed using the cancerous cervical tumor cell line HeLa, or HEK293T (kind gifts from Prof. Alfred Sze-Lok Cheng, CUHK). Cells were cultured and maintained at 37℃ with 5% $CO_2$ in Dulbecco´s modified Eagle´s medium (DMEM, Gibco-Invitrogen) supplemented with 10% fetal bovine serum (Gibco-Invitrogen) and 1% penicillin and streptomycin. Mycoplasma tests were performed routinely using MycoBlue Mycoplasma Detector (Vazyme, D101).

For plasmids, $FKBP12^{F36V}$-HA-OGT (fOGT) was obtained as previously described, or the same fOGT insert into a pLVX-puro backbone by BGI Genomes. Flag-c-Myc plasmids were purchased from MiaolingBio (P28670). pΔR8.9 and pCMV-VSV-G were kind gifts from Prof. Jingying Zhou (CUHK). $fOGT^{K852M}$ was mutated from fOGT using Mut Express MultiS Fast Mutagenesis Kit V2 (Vazyme, C216). Flag-c-$Myc^{T58A\ S415A}$ was mutated from Flag-c-Myc by Tsingke Gene. All the constructed plasmids were verified through Sanger sequencing by TsingKe Gene.

Both the forward chain (modified and unmodified) and the complementation chains of c-Myc-binding DNA (oligo) and the scrambled DNA were synthesized by Tsingke Gene (structures listed in Supplementary Figure 1). $N_3$ modification was at either 3' or 5' at the forward chain of oligo. Cy3 or biotin modification was at the 5' end of the forward chain of oligo or scrambled DNA.

Thiamet G (TMG, MedChemExpress, HY-12588), OSMI-4 (MedChemExpress, HY-114361), cycloheximide (CHX, Sigma-Aldrich, 239763-M), DBCO-$(PEG)_3$-AP1867 were dissolved in DMSO (Sigma-Aldrich, D2650) to desired concentration and applied accordingly.

#### Transient transfection

Transient transfection of plasmids or the oligo constructs (OGTAC/CBD/SD) were conducted according to the manufacturer's protocol using Lipo2000 transfection reagent (AboRo, RL0401; or Vazyme, TL-201). Taking 12-well plates as example, $1 \times 10^5$ HEK293T/HEK293T-fOGT cells were seeded into 12-well plates. For plasmids, 0.5 μg Flag-c-Myc, with or without 0.05 μg fOGT were transfected with lipo2000 (AboRo) for 24 h. For oligo constructs, OGTAC/CBD/SD was first incubated with lipo2000 (Vazyme), and then diluted in fresh medium to desired concentrations and applied to cells for 24 h.

#### Generation of the HEK293T- and HeLa-fOGT stable cell lines

HEK293T cells were used to produce lentivirus by co-transfecting pLVX-fOGT (5 μg), pΔR8.9 (5 μg), and pCMV-VSV-G (1 μg) with JetPRIME (Sartorius, 101000001). Cells were plated on 10 cm dishes at 1 × $10^6$ cells per dish and medium was replaced two hours prior to transfection with 10 mL fresh DMEM supplemented with 10% FBS. A DNA-reagent complex was prepared by combining plasmid and transfection reagent solutions, incubated at room temperature for 30 min, and added dropwise to cells; after overnight incubation the medium was replaced and viral supernatant was collected at 40 h, clarified through a 0.45 μm syringe filter, and concentrated. For generation of HEK293T-fOGT, HEK293T cells in 6-well plates were transduced with concentrated lentivirus in the presence of 8 μg/mL polybrene (MedChemExpress, HY-112735) for 16 h, medium was replaced, and 48 h later cells were selected with 2 μg/mL puromycin (MedChemExpress, HY-B1743) for 48 h; single-cell clones were isolated by limiting dilution, expanded for 10 days, and fOGT expression was confirmed by western blot, with the highest-expressing clone selected for subsequent experiments. For HeLa-fOGT generation, HeLa cells were plated at 2 × $10^5$ cells per well in 6-well plates, exposed to 1 mL viral aliquot containing 8 μg/mL polybrene for 16 h, and 48 h post-transduction medium was replaced with DMEM containing 10% FBS and 500 ng/mL G418 (ThermoFisher, 10131035). Cells were maintained under antibiotic selection for two weeks followed by monoclonal selection.

**Generation of oligo(5' or 3')-$(PEG)_3$-AP1867 and the controls**

Assembling reaction of the molecules was referred to the previous report,[1] and through a strain-promoted azide–alkyne cycloaddition (SPAAC) reaction. In detail, for the generation of oligo(5' or 3')-$(PEG)_3$-AP1867, 50 μL 100 μM forward chain of oligo(5' or 3')-$N_3$ was incubated with same amount of the complementation chain, annealing at 95℃ for 5 min and cooling to room temperature (R.T.). 20 μL 1 mM DBCO-$(PEG)_3$-AP1867 was added to the mixture, where the SPAAC reaction was performed under incubation at 37℃ with 1500 rpm shaking for 2 h. The product was further annealed once as described above. Oligo lacking the azide modification and a scrambled nucleotide chain were subjected to the same reaction conditions mixing with DBCO-$(PEG)_3$-AP1867 and were used as negative controls with the further annealing, and were used as controls - c-Myc-binding DNA complex (CBD) and scrambled DNA complex (SD). Conditions were identical to assemble the Cy3-OGTAC or Cy3-conjugated controls.

**DNA native PAGE**

To characterize and monitor the SPAAC reaction, 1 μg products from the above reactions were mixed with a 5× non-reducing loading buffer (50% glycerol, 0.25% bromophenol blue, 0.25% xylene cyanol, 10 mM Tris-HCl (pH 7.5), and 1 mM EDTA), and were subjected to 21% native PAGE (gel made from the Vazyme E305 kit). Molecular weight was indicated using the low molecular weight DNA ladder (New England Biolabs, N3233L).

**Western blot and antibodies**

For general whole cell lysate samples, $5 \times 10^5$ HeLa or HEK293T cells seeded and treated in 6-well plates were lysed using 150 μL complete RIPA-IP buffer (half-half mixed; ThermoFisher, 89900 and 87787; containing 1× protease inhibitor cocktail (TargetMol, C0001) and 1× phosphatase inhibitor cocktail (TargetMol, C0003)), collected and spun down as described above. Bicinchoninic acid (BCA; ThermoFisher, 23225) assay was performed to adjust the cell lysate to a final concentration of 2 mg/mL, and then the samples were denatured adding final 1× SDS-loading buffer (Bio-Rad, 161-0737) and heating at 95℃ for 5 min. Equal amount of protein samples for each batch were loaded for sodium dodecyl sulfate polyacrylamide gel electrophoresis (SDS-PAGE, gels made using One-Step PAGE Gel Fast Preparation Kit (8%, Vazyme, E302-01)) and blotted with corresponding antibodies, with molecular weight indicated by 180 kDa pre-stained protein markers (Vazyme, MP102-01). Analysis and quantifications of western blot results were performed using ImageJ for windows.

Primary antibodies: anti-O-GlcNAc MultiMab® (Cell Signaling Technology, 82332; 1:1000 for WB), anti-c-Myc (Santa Cruz, sc-40; 1:500 for WB), anti-HA-Tag (Vazyme, RA1004; 1:1500 for WB), anti-FlagTag (MedChemExpress, HY-P80111; 1:7000 for WB), anti-Vinculin (Santa Cruz Biotechnology or ABclonal, sc-73614 or A2752; 1:1000 or 1:50000 for WB), anti-β-actin (ABclonal, AC026 or AC004; 1:100000 or 1:3000 for WB).

Secondary antibodies: anti-rabbit HRP-linked antibody (Cell Signaling Technology, 7074; 1:8000 for WB), anti-mouse HRP-linked antibody (Cell Signaling Technology, 7076; 1:8000 for WB), anti-rabbit DyLight™ 488 antibody (ThermoFisher, 35522; 1:5000 for WB), anti-mouse DyLight™ 488 antibody (ThermoFisher, 35503; 1:5000 for WB).

**Biotin pull-down assay**

$2 \times 10^6$ HEK293T cells were seeded in a 10 cm dish. Flag-c-Myc plasmids were transfected as described above. After 24 h, cells were collected and lysed using complete RIPA-IP buffer and the concentration was adjusted to 2 mg/mL. The lysate was then divided into equal amounts, incubating at 4 ℃ overnight with biotin-oligo or unbiotinylated oligo or biotin-scrambled DNA at 0.02/0.2/2 μM, or equal amount of DMSO. All the oligonucleotides were annealed as described above. Biotin pull-down was then performed using streptavidin magnetic beads (MedChemExpress, HY-K0208) according to manufacturer's protocol and boiled in 60 μL 2× SDS-loading buffer at 95℃ for 5 min for elution. To detect pulled-down c-Myc, the eluted samples were directly applied to western blot analysis.

***In vitro* Flag-Tag pull-down assay**

5 × $10^6$ HEK293T cells were seeded in a 15 cm dish. Flag-c-Myc (10 μg) and fOGT (1 μg) plasmids were co-transfected as described above. After 24 h, cells were lysed by 1 mL RIPA-IP buffer and separated by centrifuge as described above, and divided into equal amounts. The lysate was then incubated with 10 μM TMG, or DMSO, or 1 μg/mL of both double/single-stranded CBD or oligo(5' or 3')-$(PEG)_3$-AP1867 for 2 h at R.T. Pull-down was then performed using anti-Flag magnetic beads (MedChemExpress, HY-K0207) and eluted as described above. The eluted samples were directly applied to western blot analysis.

**Immunoprecipitation (IP) and co-IP**

3 × $10^6$ HeLa-fOGT or 2 × $10^6$ HEK293T-fOGT cells were seeded in 10 cm dish for each IP reaction. Transfection of Flag-c-Myc to HEK293T-fOGT was performed at 50% - 60% confluency. After 24 h, medium was replaced by fresh warm complete DMEM containing lipo2000-oligo complexes at desired oligo concentration or lipo2000 only for 24 h. Then cells were washed by PBS and completely lysed using 600 μL complete RIPA-IP buffer. The lysate supernatant was extracted centrifuging at 14,000 RPM for 15 min at 4℃, and then applied to pre-equilibrated anti-Flag magnetic beads (MedChemExpress, HY-K0207) or anti-HA nanobody magarose beads (AlpalifeBio, KTSM1335), according to manufacturer's protocol. After gentle rotation at 4℃ overnight, the beads were washed according to manufacturer's protocol, and boiled in 80 μL 2× SDS-loading buffer at 95℃ for 5 min for elution. To detect IPed and co-IPed proteins, the eluted samples were directly applied to western blot analysis.

**Immunofluorescence (IF) staining and imaging**

5 × $10^5$ HeLa-fOGT cells were seeded into 6-well plates. Transfection of Cy3-conjugated OGTAC/CBD/SD (1 μg/mL) was performed as described above for 24 h. After removing the medium, the cells were washed with 1× Tris-Buffered Saline (TBS, containing 1× protease inhibitor cocktail)

for 5 min and fixed with pure ethanol for 15 min at -20℃. After washing, the fixed cells were blocked using 5% bovine serum albumin (BSA) dissolved in 1× TBS for 30 min, followed by anti-c-Myc primary antibody (Abcam, ab32072; 1:100 for IF) at 4℃ overnight, Alexa Fluor™ 488 secondary antibody (ThermoFisher, A-11008; 1:500 for IF) at R.T. for 1 h, and finally DAPI (ThermoFisher, 62248; 1:4000 for IF) for 15 min. Imaging was then performed using the Nikon Eclipse Ti-E fluorescence microscope and imaging system. Analysis and quantifications of IF results were performed using NIS-Elements Viewer 5.22 64-bit and ImageJ for windows.

**Cycloheximide (CHX) chase assay**

$5 \times 10^5$ HeLa-fOGT cells were seeded into 6-well plates. Transfection of OGTAC/CBD/SD (1 μg/mL) was performed as described above. After 24 h, fresh complete medium containing 50 μg/mL CHX were applied to the cells for 0, 15, 30, 45, 60 min before being lysed, normalized and denatured as mentioned above. The cell samples were then used to analyze remaining relative c-Myc level (normalized to Vinculin level) via western blot.

**Cell viability assay**

HeLa-fOGT cells (5,000 cells per well) were seeded into a 96-well plate and allowed to adhere overnight at 37 °C in a humidified incubator with 5% $CO_2$. Afterward, cells were incubated with medium containing desired concentration of OGTAC/CBD/SD for 24 h, respectively. The medium was then aspirated, and cell viability was measured using a Cell Counting Kit-8 (CCK-8, MedChemExpress, HY-K0301) assay according to the manufacturer's instructions.

**Cut&Tag-seq assay**

Cut&Tag was performed following the manufacturer's protocol (Vazyme, TD904), and next generation sequencing (NGS) of the constructed library was performed by Genewiz. In short, 2.5 $\times 10^6$ HEK293T-fOGT cells were seeded in 10 cm dishes overnight, and then Flag-c-Myc plasmids were transfected for 24 h. After changing medium, CBD or OGTAC (1 μg/mL) was treated for another 24 h. Then, 10,000 cells were collected, and subsequently incubated with ConA Beads Pro in the kit, followed by incubation with c-Myc antibody (Abcam, ab32072, 1μg every 50 μL system) at 4 ℃ overnight. Next, secondary antibody was incubated for 1 hour, and then treated with Hyperactive pA/G-Transposon Pro provided in the kit. Subsequently, DNA was extracted and amplified by PCR to construct the library, which was monitored by 1% agarose gel. Analysis of the NGS data was performed in the manufacturer's platform (Vazyme, Cut&Tag_Tool.v2.0). Data were primarily filtered in cutadapt v1.12 and normalized by spike-in and aligned to the human reference genome (hg38) in bowtie2 v2.2.9. Visualization and peak calling were performed in Macs2 v2.2.6. Difference analysis was performed in MAnorm2-utils v1.0.0 and MAnorm2 v1.2.2. Finally, a read-count threshold of 20 was applied to filter low-quality peaks. The genome-browser IGV_2.19.6 were used for visualizing c-Myc occupancy to genes individually.

**Quantitative Reverse Transcription PCR (RT-qPCR)**

$5 \times 10^5$ HeLa/HeLa-fOGT/HEK293T-fOGT cells were seeded into 6-well plates. With or without plasmids (Flag-c-Myc, and/or fOGT$^{WT}$ or fOGT$^{K852M}$) transfection as described above, the cells were treated accordingly for 24 h. Then, the cells were collected using 100 μL PBS. Total RNA was extracted using RNAfast200 (fastagen, 220010) according to the manufacturer's protocol, and was quantified using nanodrop (Monad). cDNA was synthesized from around 1 μg total RNA according to the manufacturer's protocol (Vazyme, R433). qPCR reactions (10 μL) were mixed and set according to the manufacturer's protocol (Vazyme, Q713), followed by melt-curve analysis to confirm specificity. Samples were run in technical triplicates with at least three biologically independent replicates. Relative expression was calculated by the ΔΔCt method using *HPRT1* as the reference gene and is reported as fold change relative to the SD control. Primers used were synthesized by TsingKe Gene (Supplementary Table 2).

### *Supplementary Tables.*

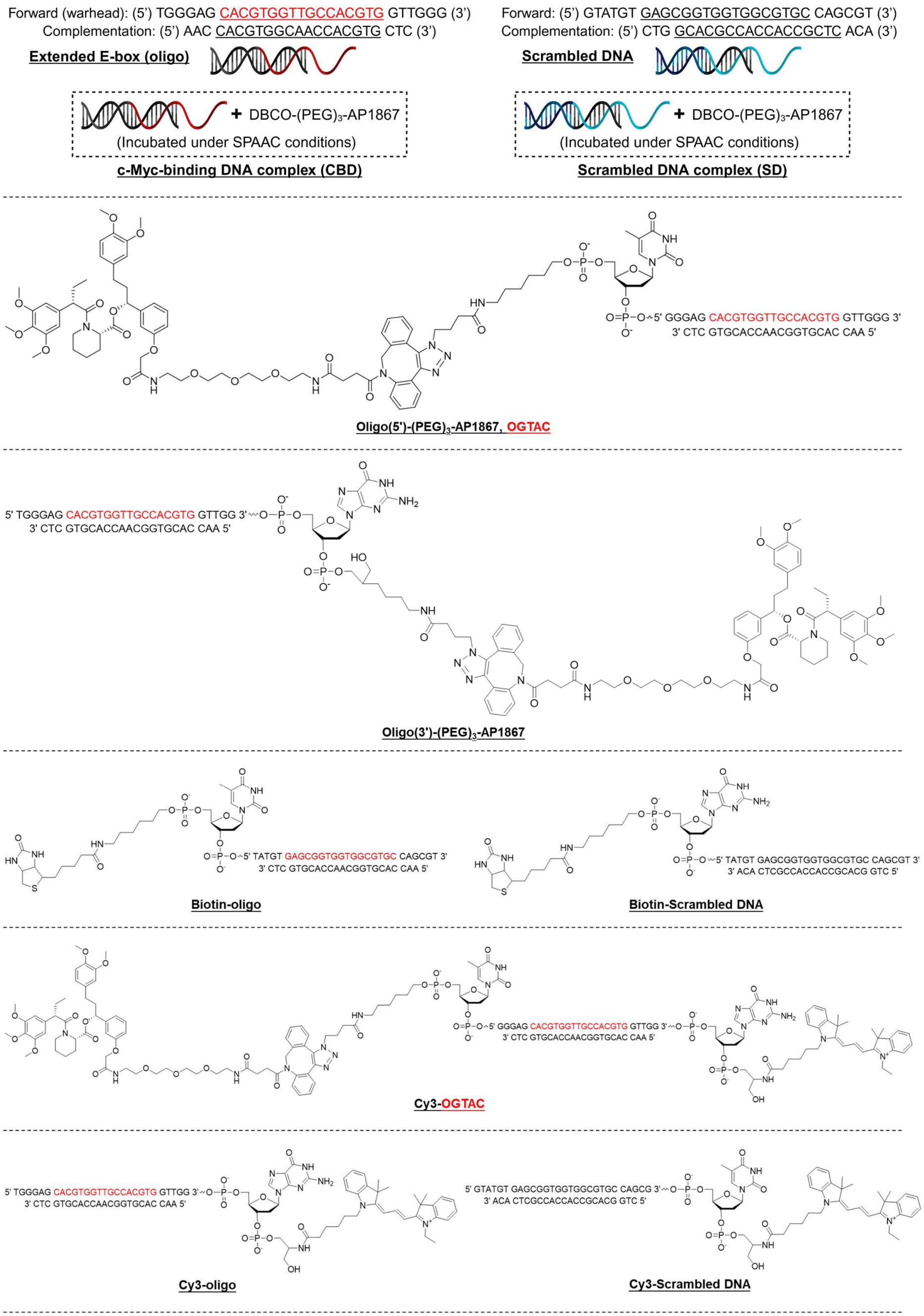

**Supplementary Table 1 Details of the molecule structures used in this study.**

| Gene | Sequence (5’ to 3’) |
|---|---|
| *MALAT1* | Forward: GGATTCCAGGAAGGAGCGAG |
| | Reverse: ATTGCCGACCTCACGGATTT |
| *MYC* | Forward: CCTGGTGCTCCATGAGGAGAC |
| | Reverse: CAGACTCTGACCTTTTGCCAGG |
| *HPRT1* | Forward: CATTATGCTGAGGATTTGGAAAGG |
| | Reverse: CTTGAGCACACAGAGGGCTACA |

**Supplementary Table 2 Primers used for qPCR in this study.**

### *Supplementary Figures.*

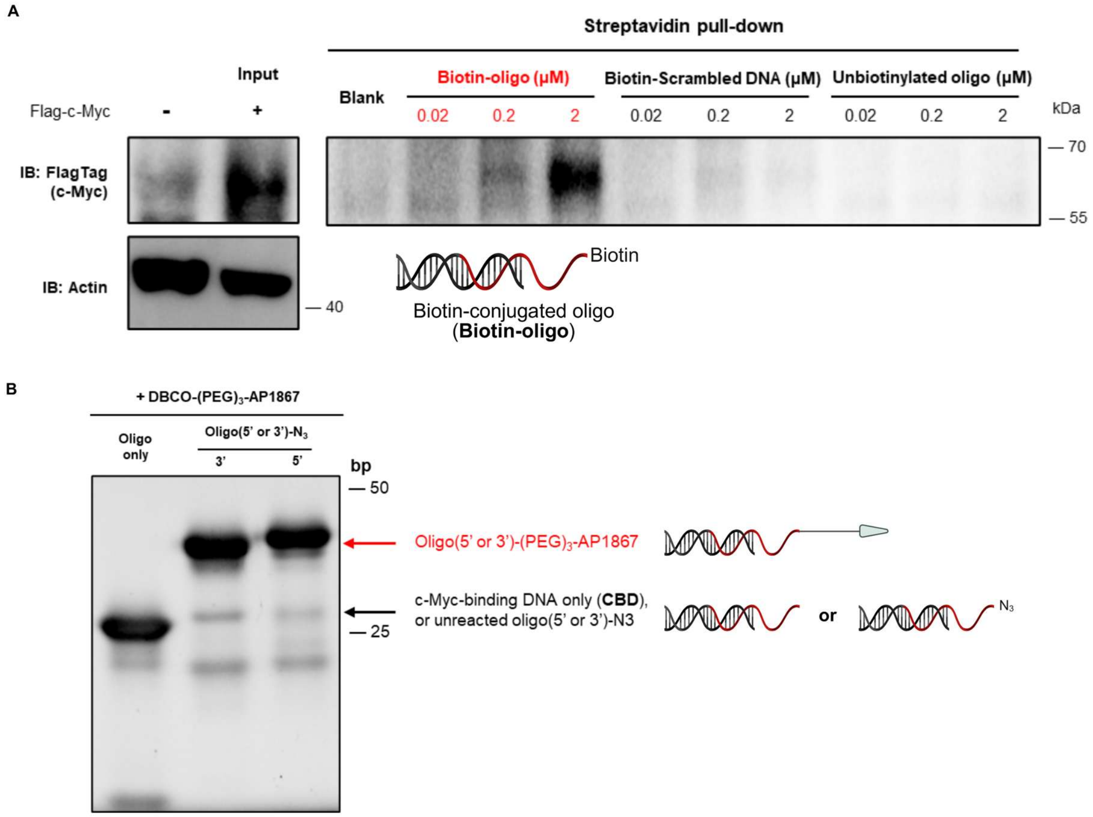

**Supplementary Figure 1 Biotin pull-down verification and native PAGE characterization.**

(A) c-Myc pull-down detected by the biotin-conjugated oligo in a dose-dependent manner, instead of the biotin-conjugated scrambled DNA or unbiotinylated oligo. Indicated concentrations of biotin-oligo, biotin-scrambled DNA, or unbiotinylated oligo were incubated with the lysate of HeLa (2 mg/mL) overexpressing Flag-c-Myc, and then with streptavidin magnetic beads for 1 h before elution. (B) Incorporation of DBCO-$(PEG)_3$-AP1867 onto the oligo(5' or 3')-$N_3$ led to an increase of molecular weight (1155 Da), shown clearly by the 21% native PAGE.

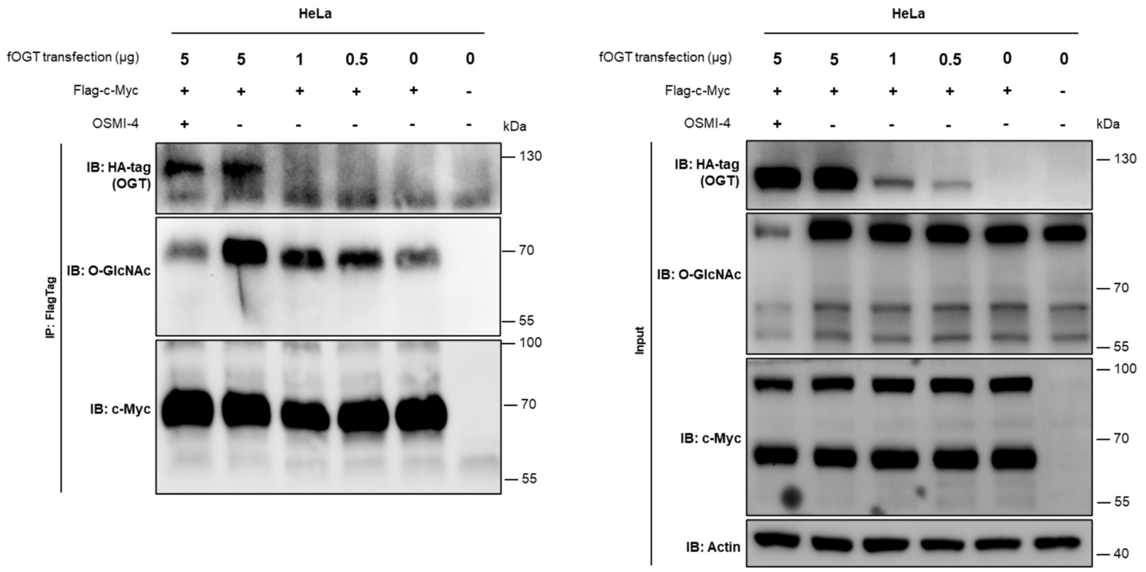

**Supplementary Figure 2 Detection of c-Myc O-GlcNAcylation.**

Detected O-GlcNAcylation of IPed c-Myc by MultiMab O-GlcNAcylation antibody. Flag-c-Myc and different amounts of fOGT were overexpressed in HeLa cells for 24 h, with/without the subsequent OSMI-4 treatment (10 μM).

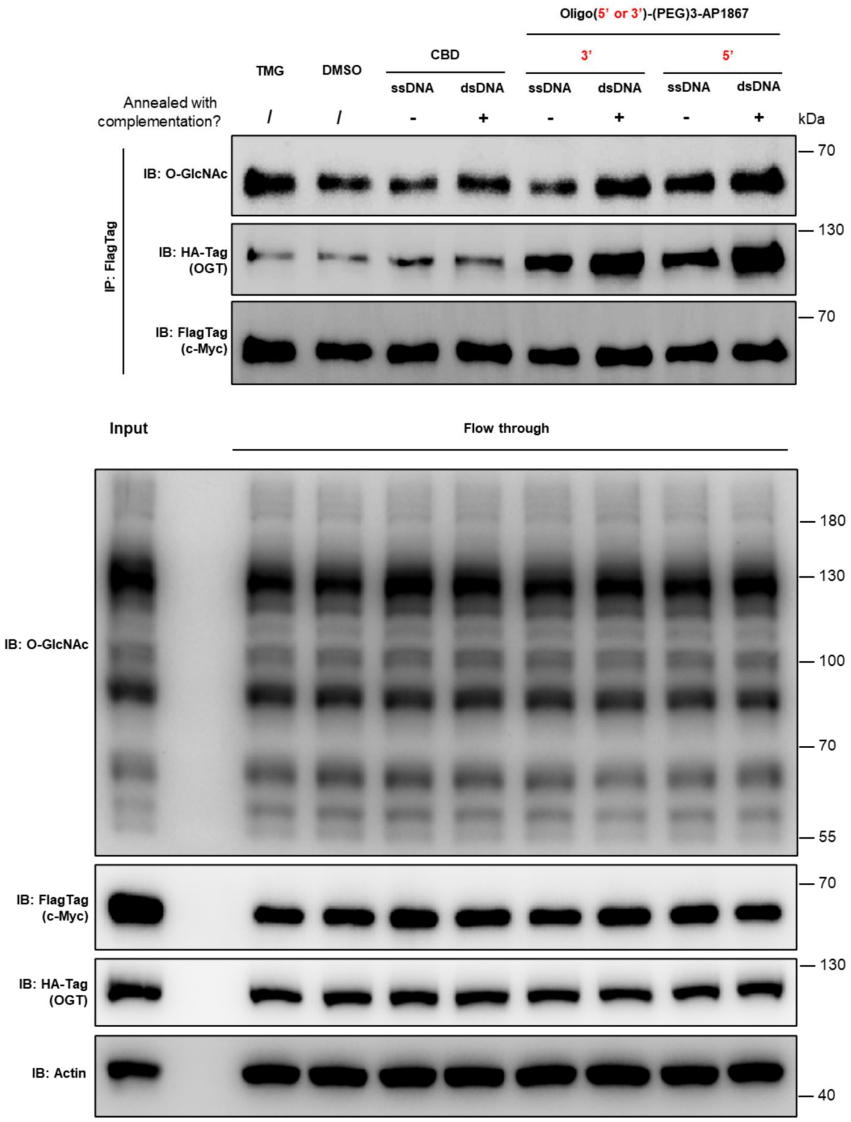

**Supplementary Figure 3 Induction effects of oligo(5' or 3')-(PEG)$_3$-AP1867 in lysate.**

Induced c-Myc O-GlcNAcylation and c-Myc-fOGT co-IP by the bifunctional oligo(5' or 3')-(PEG)$_3$-AP1867 constructs in lysate, where both annealed or unannealed oligos were tested. The constructs (1 μg/mL) or TMG (10 μM) or DMSO were incubated with the lysate of HEK293T-fOGT (2 mg/mL) overexpressing Flag-c-Myc for 2 h at R.T., followed by IP using the FlagTag magnetic beads.

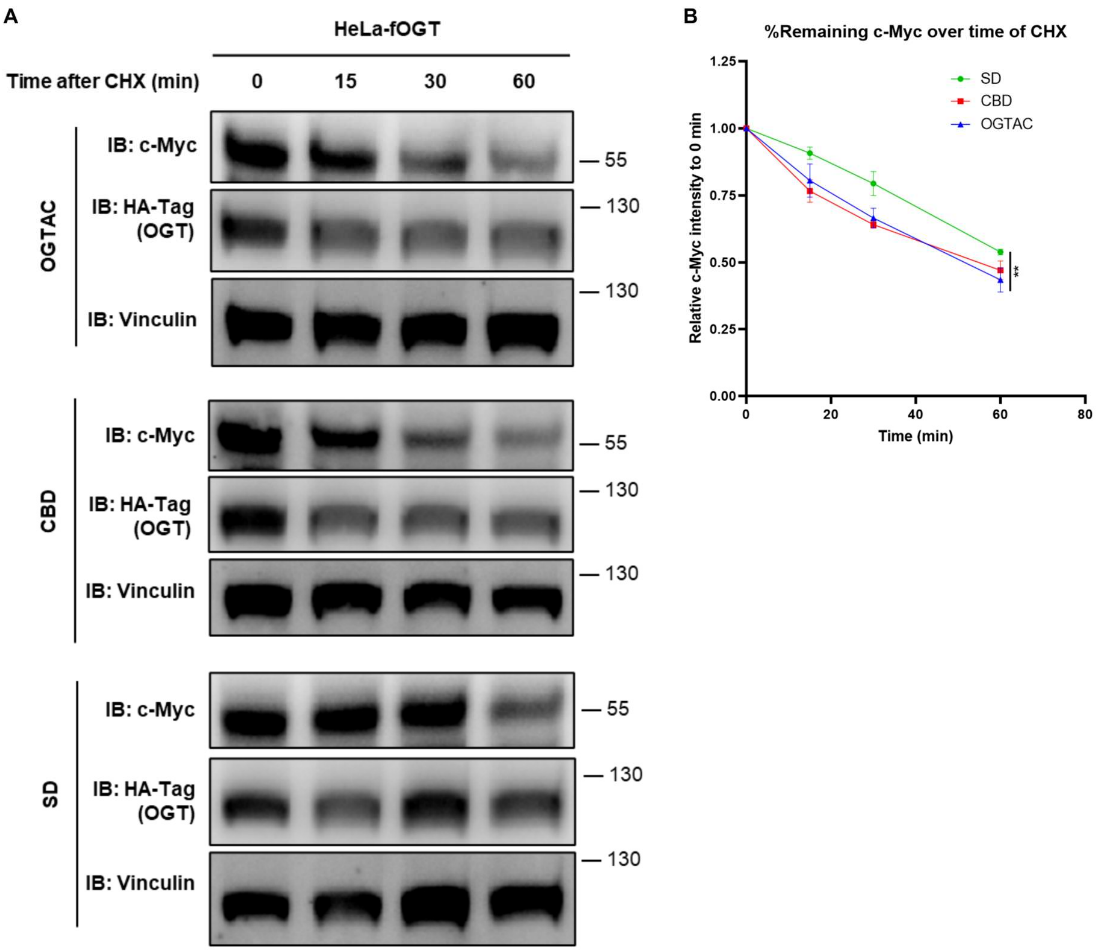

**Supplementary Figure 4 CHX chase assay to determine c-Myc stability upon treatments.**

(A) CHX chase assay to determine c-Myc stability upon treatment of OGTAC/CBD/SD (1 μg/mL, 24 h). HeLa-fOGT cells were treated accordingly, and collected to analyze remaining c-Myc level at 0/15/30/60 min after CHX treatment (50 μg/mL). (B) Quantification of relative endogenous c-Myc level (c-Myc normalized to vinculin) fold change, normalized to the 0 min level. Error bars represent the mean (SD) from *n* = 3 biologically independent experiments. Statistical significance for each quantification was assessed using a two-way ANOVA. ***p* < 0.01; ns, not significant.

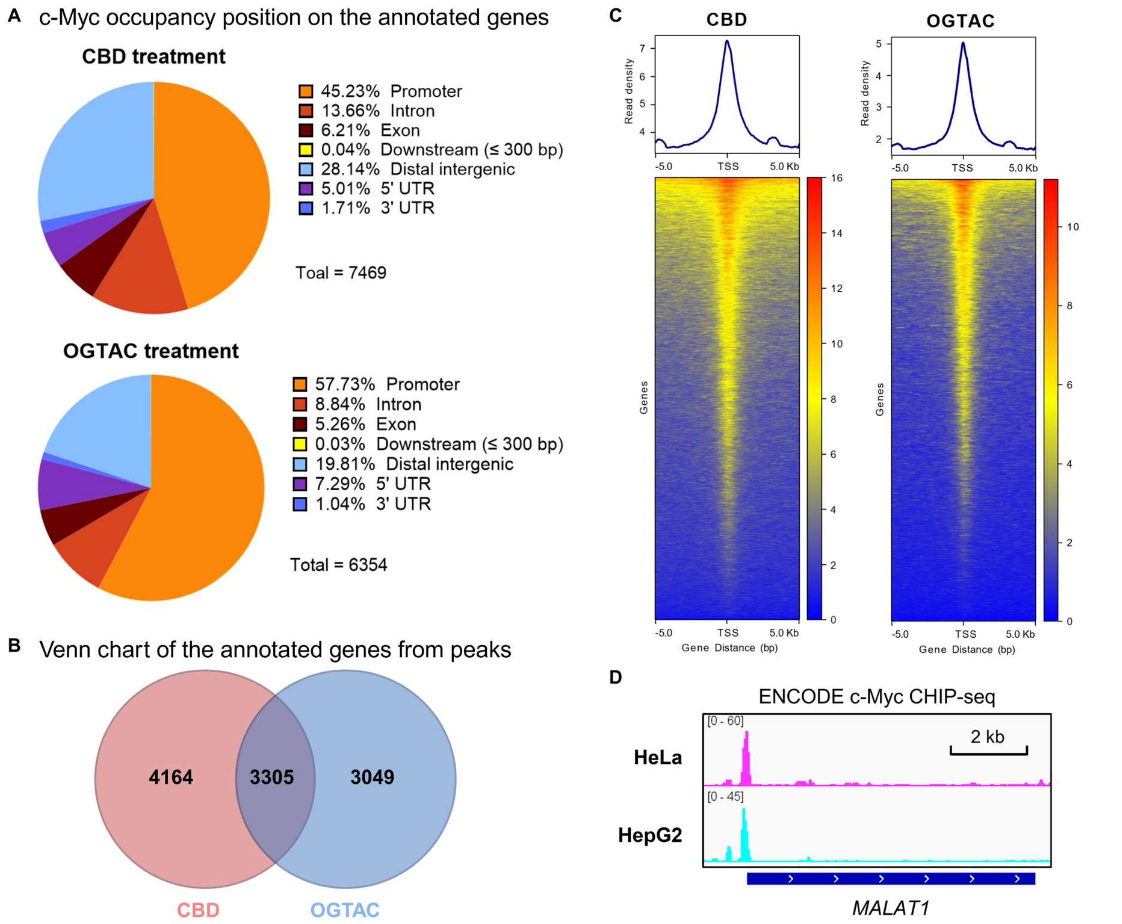

**Supplementary Figure 5 c-Myc occupancy to genome (Cut&Tag-seq) and *MALAT1* (ENCODE CHIP-seq).**

(A) Compared to CBD, OGTAC changed the overall c-Myc-occupancy position of genes. Specifically, c-Myc bound less to the distal intergenic area (including enhancers) and more at the promoter area in the OGTAC group. (B) For the genes annotated from the captured DNA fragments in Cut&Tag, OGTAC treatment led to a different set of genes being bound to c-Myc compared to CBD. (C) Heatmap of read signal enrichments across a ± 5 kb window centered on peak summits from CUT&Tag-seq showing c-Myc occupancy at the genome. X-axis represents distance from the transcription starting sites (TSS), and y-axis represents signal intensity (larger values indicate stronger occupancy). OGTAC treatment showed weakening overall c-Myc genomic occupancy compared to CBD. Reads from next-generation sequencing were normalized by spike-in. (D) ENCODE c-Myc ChIP-seq showing c-Myc occupancy at *MALAT1* promoter in HeLa cell line (magenta; accession: ENCFF375REM) and HepG2 (cyan; accession: ENCFF388VAV).[2]

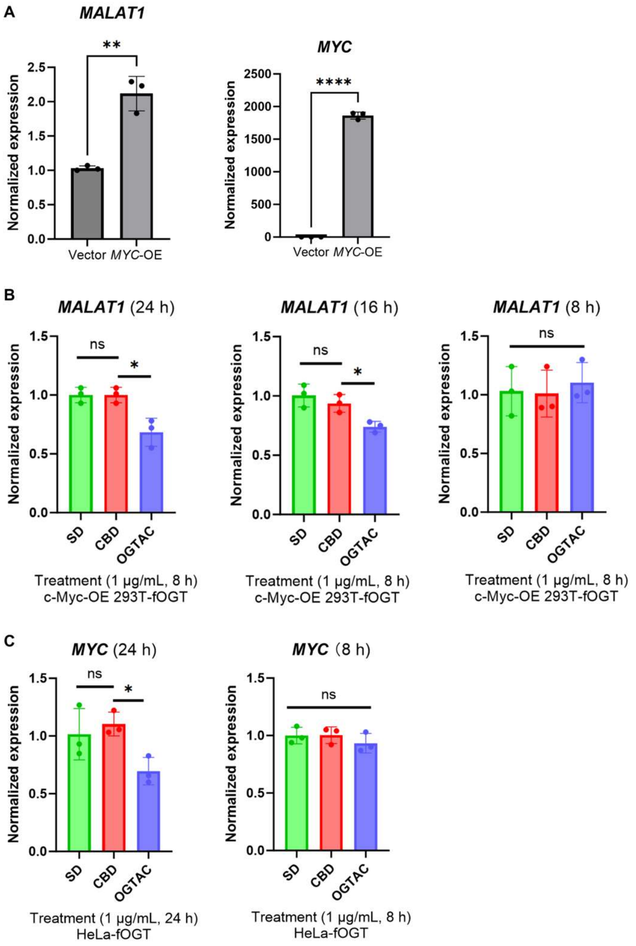

**Supplementary Figure 6 *MALAT1* and *MYC* expression in HEK293T-fOGT or HeLa-fOGT upon treatments.**

(A) Normalized expression of *MALAT1* in HEK293T cells with or without overexpressed c-Myc.

(B) Normalized expression of *MALAT1* in HEK293T-fOGT cells after 24 h, 16 h, or 8 h treatment with 1 μg/mL OGTAC, expressed relative to SD and CBD controls.

(C) Normalized expression of *MYC* in HeLa-fOGT cells after 24 h or 8 h treatment with 1 μg/mL OGTAC, expressed relative to SD and CBD controls.

For all panels, error bars represent the mean (SD) from $n = 3$ biologically independent experiments, where each point represents the mean of $n = 3$ technical replicates. Relative expression was calculated as target gene/HPRT1 and is presented as fold change relative to the SD control. Statistical significance for the quantifications was determined using an unpaired Student's $t$-test. *$p < 0.05$; **$p < 0.01$; ****$p < 0.0001$; ns, not significant.

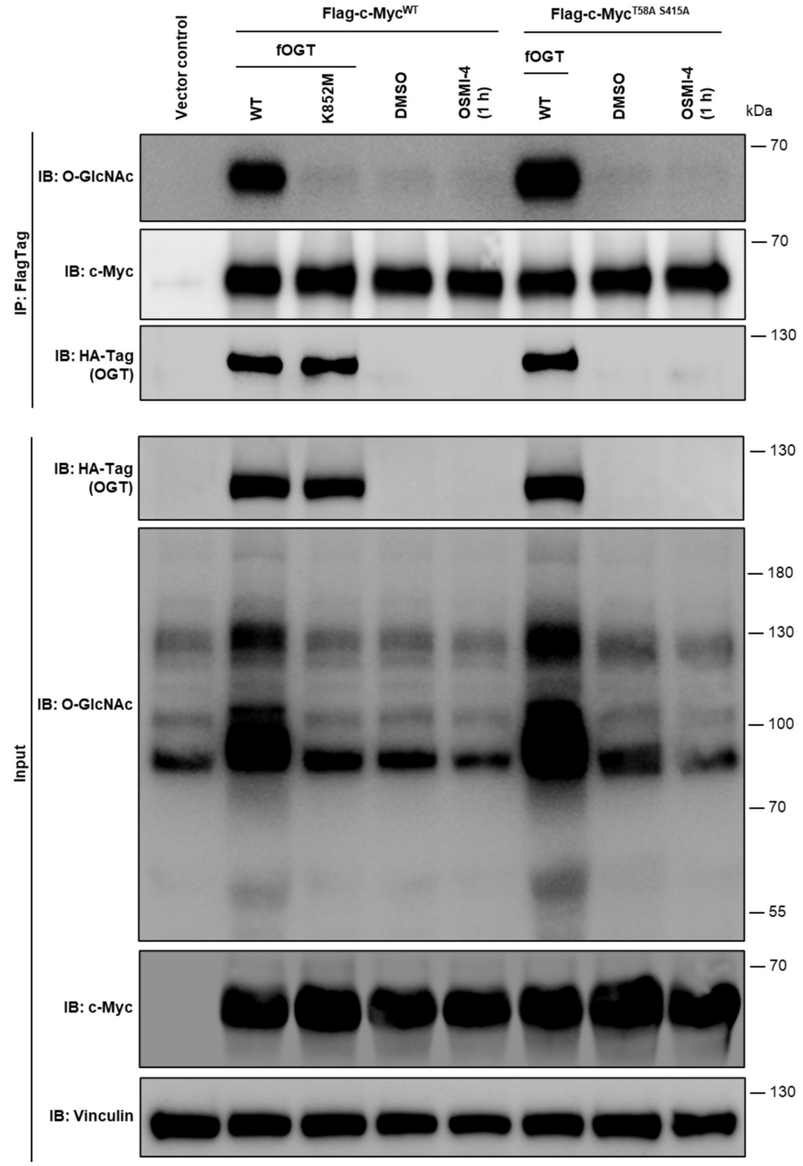

**Supplementary Figure 7 O-GlcNAcylation of c-Myc (WT or T58A S415A) with/without fOGT (WT or K852M).**

O-GlcNAcylation of c-Myc (WT or T58A S415A) was determined upon the overexpression of fOGT$^{WT}$ or fOGT$^{K852M}$ or vector. Compared to fOGT$^{WT}$, fOGT$^{K852M}$ lost the O-GlcNAcylation catalytic activity on both c-Myc and the proteome, while it retained normal interaction with c-Myc. Interestingly, c-Myc with the previously reported O-GlcNAcylation sites[3, 4] mutated (c-Myc$^{T58A\ S415A}$) still showed increased O-GlcNAcylation upon fOGT$^{WT}$ overexpression.

### *Synthesis method of DBCO-(PEG)$_3$-AP1867.*

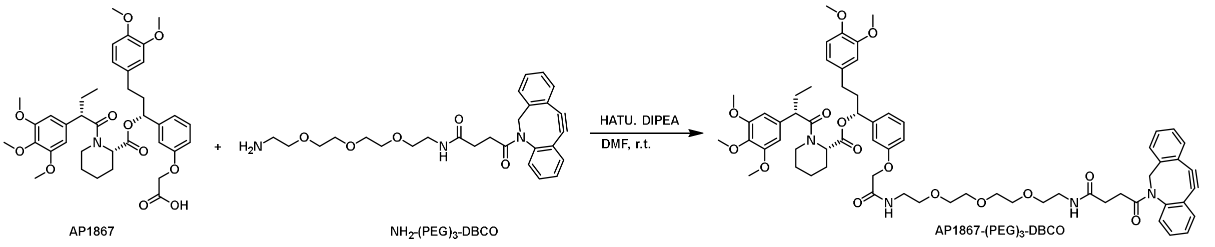

**Supplementary Scheme 1 The synthesis route of DBCO-(PEG)$_3$-AP1867.**

All solvents were anhydrous quality purchased from Meryer. Unless otherwise noted, chemical starting materials are purchased from BidePharm without further purification. Spectra were acquired on Bruker spectrometers: $^{1}$H NMR: 400 or 700 MHz (recorded at 400/700 MHz); $^{13}$C NMR: 176 MHz (recorded at 176 MHz). Chemical shifts are reported in parts-per million (ppm) relative to tetramethylsilane. Spectra were referenced according to the solvent residual peak (CDCl3 7.26 ppm, 77.0 ppm). Reactions were monitored by thin layer chromatography and the products were purified using fast column chromatography. Mass spectrometry was performed on Agilent LC-MS/MS system consisted of two Agilent 1290 series pumps and auto-sampler, coupled with 6430 triple quadrupole mass spectrometer equipped with ESI source (Agilent Technologies, Inc., Santa Clara, CA, USA).

Commercially available AP1867 (25.0 mg, 0.036 mmol, 1.0 eq, Bide Pharm) was dissolved in anhydrous DMF (1 mL). To this solution were added HATU (20.5 mg, 0.054 mmol, 1.5 eq) and DIPEA (13.9 mg, 0.108 mmol, 3.0 eq). The reaction mixture was stirred at room temperature for 0.5 h, followed by the addition of commercially available $NH_2$-(PEG)$_3$-DBCO (17.2 mg, 0.036 mmol, 1.0 eq, Bide Pharm). After stirring overnight at room temperature, the mixture was extracted with ethyl acetate (3 × 10 mL). The combined organic layers were washed with saturated brine, dried over anhydrous $Na_2SO_4$, filtered, and concentrated under reduced pressure. The residue was adsorbed onto silica gel and purified by flash column chromatography (eluent: DCM/MeOH from 30:1 to 15:1, v/v) to afford the target product as a yellow oily solid (28.4 mg, 68.3% yield).

$^{1}$H NMR (700 MHz, $CDCl_3$) δ 7.65 (d, $J$ = 7.5 Hz, 1H), 7.53 – 7.49 (m, 1H), 7.40 – 7.38 (m, 1H), 7.37 – 7.36 (m, 2H), 7.35 – 7.30 (m, 1H), 7.30 – 7.27 (m, 1H), 7.23 (d, $J$ = 7.5 Hz, 1H), 7.19 – 7.16 (m, 1H), 7.12 – 7.09 (m, 1H), 6.79 – 6.75 (m, 3H), 6.68 – 6.62 (m, 3H), 6.41 (d, $J$ = 2.1 Hz, 2H),

6.20 (s, 1H), 5.62 (dd, $J$ = 8.2, 5.5 Hz, 1H), 5.47 – 5.44 (m, 1H), 5.13 (dd, $J$ = 13.9, 2.8 Hz, 1H), 4.49 – 4.46 (m, 2H), 3.85 – 3.84 (m, 6H), 3.83 (s, 2H), 3.78 (s, 2H), 3.68 (s, 5H), 3.66 – 3.63 (m, 1H), 3.60 – 3.59 (m, 4H), 3.58 – 3.56 (m, 4H), 3.56 – 3.54 (m, 2H), 3.53 – 3.50 (m, 2H), 3.48 (s, 2H), 3.45 – 3.38 (m, 2H), 3.33 – 3.30 (m, 2H), 2.84 – 2.75 (m, 2H), 2.57 – 2.52 (m, 1H), 2.48 – 2.42 (m, 2H), 2.33 – 2.28 (m, 1H), 2.18 – 2.13 (m, 1H), 2.10 – 2.04 (m, 2H), 1.96 – 1.89 (m, 2H), 1.74 – 1.67 (m, 4H), 1.63 – 1.54 (m, 2H), 1.43 (m, $J$ = 13.0, 9.1, 4.0 Hz, 2H), 1.13 (d, $J$ = 6.5 Hz, 1H).

$^{13}$C NMR (176 MHz, $CDCl_3$) δ 172.73, 172.39, 172.19, 170.68, 168.32, 157.39, 153.28, 151.49, 149.01, 148.19, 147.40, 142.38, 136.63, 135.45, 133.41, 132.32, 129.95, 129.51, 128.78, 128.26, 127.83, 127.15, 125.60, 123.30, 122.54, 120.27, 119.82, 114.77, 114.05, 112.94, 111.70, 111.28, 108.01, 104.98, 104.56, 75.76, 70.56, 70.35, 70.27, 69.85, 69.79, 67.45, 60.90, 56.38, 56.06, 56.01, 55.93, 55.61, 52.14, 51.01, 50.88, 43.58, 39.24, 38.88, 38.40, 31.38, 31.24, 30.22, 29.83, 28.47, 26.91, 25.44, 23.61, 21.06, 12.70.

HRMS: calcd for $C_{65}H_{78}N_4O_{15}$ $[M+Na]^+$: 1177.53559, Found: 1177.53496.

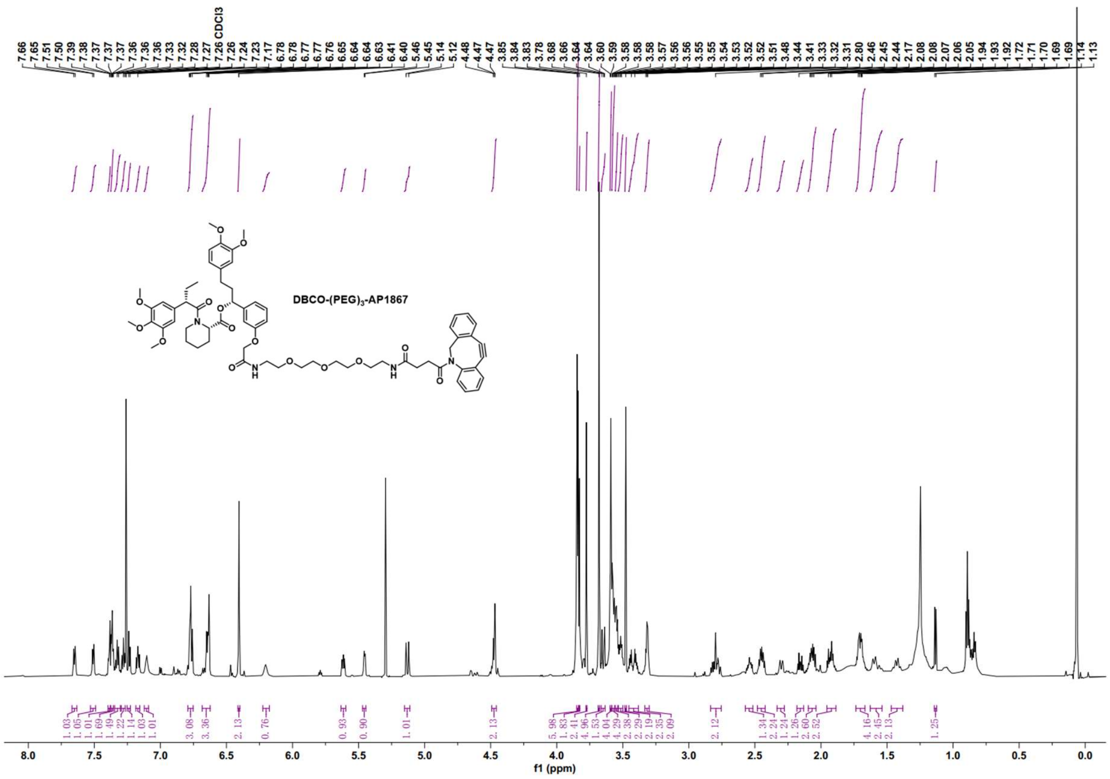
DBCO-(PEG)3-AP1867
f1 (ppm)

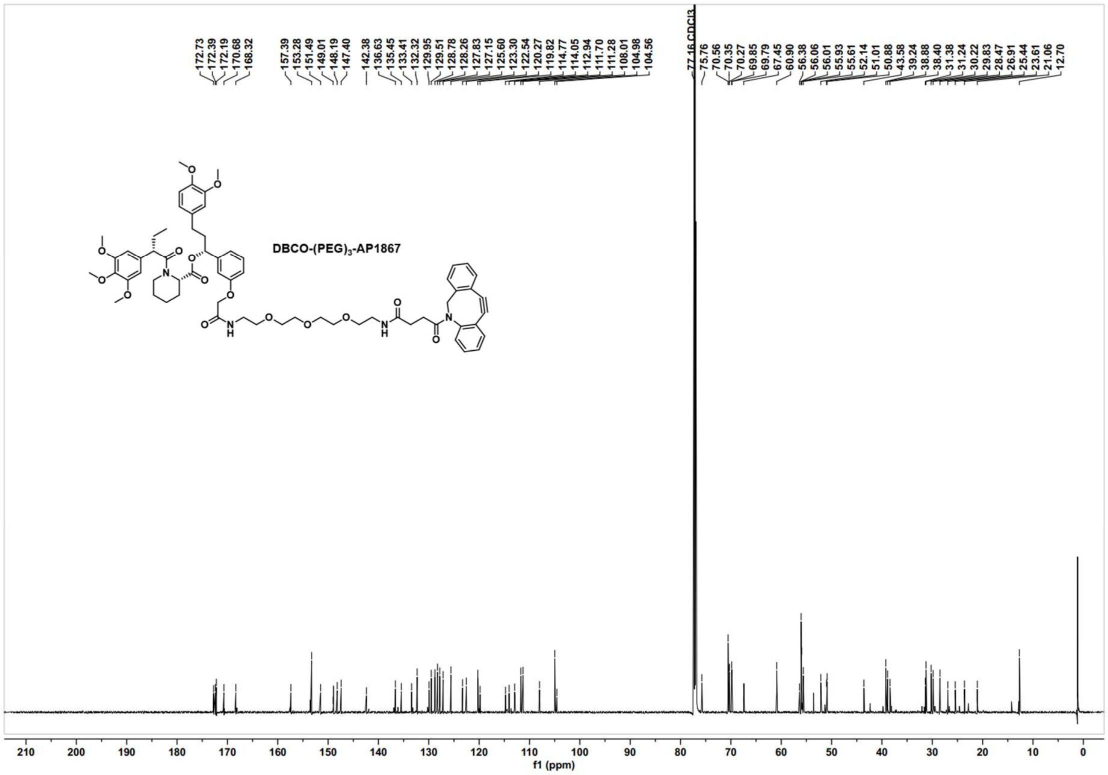
DBCO-(PEG)3-AP1867
f1 (ppm)

### ***References.***

1. Samarasinghe, K. T. G.; An, E.; Genuth, M. A.; Chu, L.; Holley, S. A.; Crews, C. M., OligoTRAFTACs: A generalizable method for transcription factor degradation. *RSC Chemical Biology* **2022,** *3* (9), 1144-1153.

2. Landt, S. G.; Marinov, G. K.; Kundaje, A.; Kheradpour, P.; Pauli, F.; Batzoglou, S.; Bernstein, B. E.; Bickel, P.; Brown, J. B.; Cayting, P.; Chen, Y.; DeSalvo, G.; Epstein, C.; Fisher-Aylor, K. I.; Euskirchen, G.; Gerstein, M.; Gertz, J.; Hartemink, A. J.; Hoffman, M. M.; Iyer, V. R.; Jung, Y. L.; Karmakar, S.; Kellis, M.; Kharchenko, P. V.; Li, Q.; Liu, T.; Liu, X. S.; Ma, L.; Milosavljevic, A.; Myers, R. M.; Park, P. J.; Pazin, M. J.; Perry, M. D.; Raha, D.; Reddy, T. E.; Rozowsky, J.; Shoresh, N.; Sidow, A.; Slattery, M.; Stamatoyannopoulos, J. A.; Tolstorukov, M. Y.; White, K. P.; Xi, S.; Farnham, P. J.; Lieb, J. D.; Wold, B. J.; Snyder, M., ChIP-seq guidelines and practices of the ENCODE and modENCODE consortia. *Genome Res* **2012,** *22* (9), 1813-31.

3. Chou, T.-Y.; Hart, G. W.; Dang, C. V., c-Myc Is Glycosylated at Threonine 58, a Known Phosphorylation Site and a Mutational Hot Spot in Lymphomas. *Journal of Biological Chemistry* **1995,** *270* (32), 18961-18965.

4. Wang, H.; Sun, J.; Sun, H.; Wang, Y.; Lin, B.; Wu, L.; Qin, W.; Zhu, Q.; Yi, W., The OGT–c-Myc–PDK2 axis rewires the TCA cycle and promotes colorectal tumor growth. *Cell Death & Differentiation* **2024,** *31* (9), 1157-1169.
